# Prediction of the closed conformation and insights into the mechanism of the membrane enzyme LpxR

**DOI:** 10.1101/309096

**Authors:** Graham M Saunders, Hannah E Bruce Macdonald, Jonathan W Essex, Syma Khalid

## Abstract

Covalent modification of outer membrane lipids of Gram-negative bacteria can impact the ability of the bacterium to develop resistance to antibiotics as well as modulating the immune response of the host. The enzyme LpxR from *Salmonella typhimurium* is known to deacylate lipopolysaccharide molecules of the outer membrane, however the mechanism of action is unknown. Here we employ Molecular Dynamics and Monte Carlo simulations to study the conformational dynamics and substrate binding of LpxR in representative outer membrane models and also detergent micelles. We examine the roles of conserved residues and provide an understanding of how LpxR binds its substrate. Our simulations predict that the catalytic H122 must be Nε-protonated for a single water molecule to occupy the space between it and the scissile bond, with a free binding energy of -8.5 kcal mol^-1^. Furthermore, simulations of the protein within a micelle enable us to predict the structure of the putative ‘closed’ protein. Our results highlight the need for including dynamics, a representative environment and the consideration of multiple tautomeric and rotameric states of key residues in mechanistic studies; static structures alone do not tell the full story.

## INTRODUCTION

The outer membrane (OM) of Gram-negative bacteria is a formidable barrier to the permeation of molecular species seeking to enter the bacterial cell^1^. It is only selectively permeable, enabling molecules essential for the survival of the bacteria such as nutrients to get across the membrane, but excluding those that are harmful, such as antibacterial agents. The chemical natures of the lipids of the membrane are thought to play a key role in achieving this selective permeability. The membrane is a lipid bilayer, with the leaflet facing the external environment, known as the outer leaflet, almost exclusively composed of lipopolysaccharide molecules, whereas the inner leaflet contains a mixture of zwitterionic and anionic lipids. To date, lipopolysaccharide is perhaps the most chemically complex natural lipid known. The internal membrane component is known as lipid A, this is covalently linked to a polymer of sugars, some of which are phosphorylated. The chemical structure of the lipopolysaccharide molecules varies across bacterial species and sometimes even within one species, for example in *E. coli* and *S. typhimurium* the lipid A component has six acyl tails, whereas in *P. aeruginosa* it only has five tails^2^. In *E. coli* and *S. typhimurium*, these are laurate and myristate tails in a ratio of 5:1. LPS may be referred to as smooth, rough or deep rough LPS, referring to the number of sugars attached to the lipid A segment. Smooth LPS includes the O-antigen and the full core segment, rough just the six core sugars and deep rough only two keto-deoxyoctulosonate (Kdo) sugars. Rough and deep rough LPS are also referred to as Ra and Re LPS respectively.

A number of pathogenic bacteria have been shown to synthesize LPS molecules with modified lipid A. Some of these modifications to lipid A can facilitate the development of resistance to drugs, for example addition of the L-Ara4N group which is positively charged at pH 7, neutralizes the negative charge of a lipid A phosphate group in *E. coli, S. typhimurium* and *P. aeruginosa*, which reduces the susceptibility of these bacteria to antimicrobial peptides. A number of different bacterial enzymes that catalyse the covalent modification of LPS have been identified. Three of these enzymes, all of which modify lipid A tails, PagP, PagL, and LpxR, are embedded in the outer membrane^3–5^. Positioning of coarse-grain and united-atom LpxR models with respect to phosphate headgroups can be seen in Figures 1a and 1b. LpxR catalyses the removal of two acyl chains of lipid A, in the form of 3-(tetradecanoyloxy)tetradecanoic acid (Figure 1c). The X-ray structure of the protein has been resolved to 1.9 Å resolution, and while the structure of the lipid substrate bound to the protein remains elusive, mutational studies have identified residues that are essential for catalytic activity thus providing clues to the location of the active site (Figure 1d)^5^. The X-ray structure of LpxR was obtained by co-crystallizing the protein with Zn^2+^, and while there is some evidence of Zn^2+^ binding to LpxR in the X-ray structure, the density is low and likely indicative of partial occupation. Interestingly, it has been shown that while Ca^2+^ is essential for LpxR activity, some other divalent cations, such as Sr^2+^ and Cd^2+^, but not Zn^2+^+ can replace Ca^2+^ without total loss of catalytic activity^5^. Thus, while we know Ca^2+^ is essential, it is unclear from the X-ray structure precisely where it is bound for catalytic activity to occur. Based on their structures and mechanistic predictions, the mechanism of deacylation in LpxR is speculated to be similar to that displayed by a number of phospholipase A2 enzymes^6,7^. The proposed mechanism of deacylation is thought to occur via a histidine, H122 acting as a base by activating a nearby water molecule, so the latter can react with the substrate lipid molecule to hydrolyse the ester bond. In the X-ray structure there is a water molecule resolved near H122, which may well be the crucial mechanistic water. While the structure of LpxR, the location of the water and the docked lipid substrate provide a plausible static model for the reaction mechanism of the enzyme, the dynamic stability of the model protein-substrate complex, the location of any additional key water and ion binding sites (especially given the very low electron density for Zn^2+^ in the X-ray structure), and the effect of the complex on the local membrane are still unexplored and thus the mechanism of action has not been proven.

**Figure 1.**
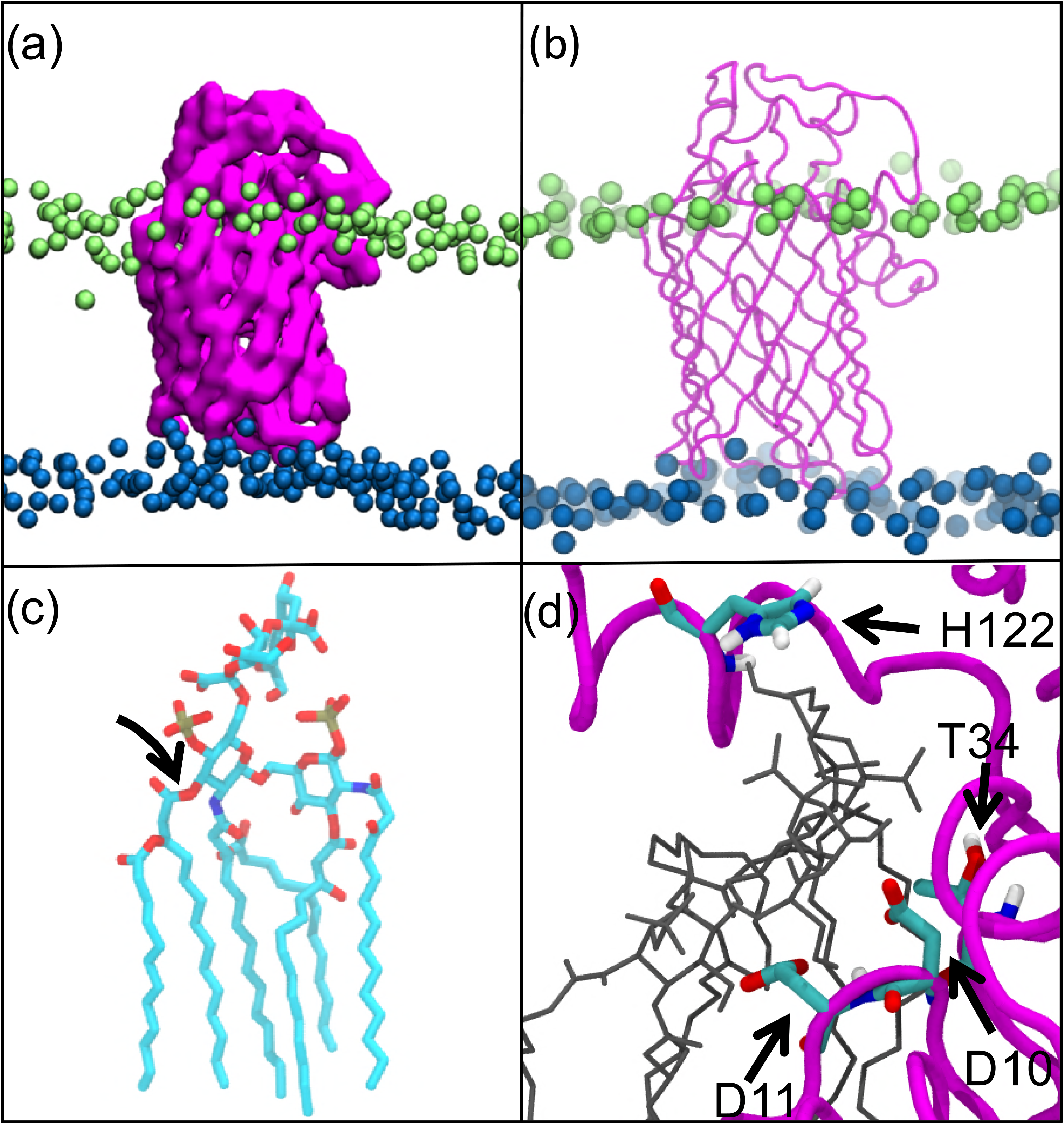
(a) coarse-grain and (b) united atom models of LpxR from the X-ray structure (PDB code 3FID) embedded within a model of the OM. Ra LPS headgroups are shown in lime, phospholipid headgroups are shown in blue, lipids tails are omitted for clarity. (c) Re LPS with scissile bond indicated and hydrogen atoms removed for clarity. (d) Close-up of the docked Re LPS molecule (black) with residues thought to be key for catalysis through mutagenesis studies, highlighted^5^.

To this end, here we present a molecular simulation study in which we use a combination of Molecular Dynamics and Monte Carlo simulations to investigate the molecular interactions between the LpxR and the Re LPS substrate, with the specific aim of uncovering mechanistic insights into the process of deacylation. Details of the calculations are presented at the end of the paper, but for ease of reading we present the tables summarising the Molecular Dynamics (Table 1) and Monte Carlo (Table 2) simulations below.

**Table 1.** Summary of all MD simulations reported in this work.

| System | Notes | Length of each simulation | Temperature and number of repeats |
| --- | --- | --- | --- |
| Apo Model | X-ray structure with water and $\text{Ca}^{2+}$ as reported by Rutten et al. | 300 ns | 310 K (x2)<br>323 K (x1) |
| Apo Model* | As above, but with positional restraints for first 100 ns | 300 ns | 310 K (x2)<br>323 K (x1) |
| Apo Unbiased | X-ray structure with key water and $\text{Ca}^{2+}$ removed from starting structure | 300 ns | 310 K (x2)<br>323 K (x1) |
| Ligand-Bound Model | X-ray structure with water and $\text{Ca}^{2+}$ and modelled in substrate as reported by Rutten et al<sup>5</sup> | 300 ns | 310 K (x2)<br>323 K (x1) |
| Ligand-Bound Model* | As above, but with positional restraints for first 100 ns | 300 ns | 310 K (x2)<br>323 K (x1) |
| Ligand-Bound Unbiased | X-ray structure with modelled in substrate, but without water and $\text{Ca}^{2+}$ | 300 ns | 310 K (x2)<br>323 K (x1) |
| H122_p** | H122 is Nε-protonated | 300 ns | 310 K (x1) |
|  |  |  | 323 K (x1) |
| D10A | Single point mutation in protein | 300 ns | 310 K (x2)<br>323 K (x1) |
| D11A | Single point mutation in protein | 300 ns | 310 K (x2)<br>323 K (x1) |
| T34A | Single point mutation in protein | 300 ns | 310 K (x2)<br>323 K (x1) |
| H122A | Single point mutation in protein | 300 ns | 310 K (x2)<br>323 K (x1) |
| Bond Break Model | Scissile bond removed in substrate | 500 ns | 310 K (x2)<br>323 K (x1) |
| Bond Break Unbiased | As above but with key water and $\text{Ca}^{2+}$ removed from starting structure | 500 ns | 310 K (x2)<br>323 K (x1) |
| Micelle | Protein in self-assembled DPC micelle | 500 ns | 323 K (x3) |
| CG_LpxR | Coarse-grain system | 2 $\mu\text{s}$ | 310 K (x2)<br>323 K (x1) |
| **Total Simulation Time: 20.1 $\mu\text{s}$** |  |  |  |

\* In these systems the protein, $\text{Ca}^{2+}$ ion and modelled water molecule were subjected to positional restraints for the first 100 ns of simulation, see the methods section for more details.
\*\* H122 is $\text{N}_\epsilon$ -protonated in this simulation, whereas it is $\text{N}_\delta$ in all other simulations.

* In these systems the protein, Ca^2+^ ion and modelled water molecule were subjected to positional restraints for the first 100 ns of simulation, see the methods section for more details. ** H122 is Nε-protonated in this simulation, whereas it is Nδ□in all other simulations.

**Table 2.** Summary of all GCMC simulations reported in this work.

| Simulation | Repeats | B-value(s) | Production Steps | GCMC Box |
| --- | --- | --- | --- | --- |
| H122ε | 3 | -40.79 to -10.79 | 20 M | 0.2 x 0.2 x 0.2 nm<sup>3</sup> |
| H122δ | 3 | -40.79 to -10.79 | 20 M | 0.2 x 0.2 x 0.2 nm<sup>3</sup> |
| H122$\epsilon^*$ | 3 | -7.94 | 40 M | 0.73 x 0.49 x 1.16 nm<sup>3</sup> |
| H122$\delta^*$ | 3 | -7.94 | 40 M | 0.73 x 0.49 x 1.16 nm<sup>3</sup> |
| H122$\epsilon^{**}$ | 3 | -40.79 to -10.79 | 20 M | 0.2 x 0.2 x 0.2 nm<sup>3</sup> |
| H122$\delta^{**}$ | 3 | -40.79 to -10.79 | 20 M | 0.2 x 0.2 x 0.2 nm<sup>3</sup> |

\*\* In these systems, the imidazole ring of H122 was flipped 180°.

* These systems refer to the water location identification simulations. ** In these systems, the imidazole ring of H122 was flipped 180°.

## RESULTS AND DISCUSSION

### Protein orientation and stability

Coarse-grain simulations were performed to provide an unbiased prediction of the localisation and orientation of LpxR with respect to the outer membrane. The densities of molecular components as well tilt angle of the protein with respect to the plane of the membrane calculated from the coarse-grain simulations were used to set up the atomistic systems. Comparisons of these metrics between atomistic and coarse-grain simulations are provided in the SI. The principal axis of the protein barrel remains tilted by ~13° relative to the bilayer normal during simulations at both resolutions. As has been reported for other outer membrane proteins, orientation of the protein is maintained in part by the aromatic residues on the outer surface of the barrel being positioned at the lipid headgroup-tail interface of the membrane^8^. Interestingly we observe some local deformations of the membrane: specifically, the membrane is ~0.8 nm thinner within a radius of ~1.2 nm around the protein, compared to further away in what can be considered the bulk lipid region (Figure S1).

Comparison of the order parameters of LPS acyl tails within 0.5 nm of the protein and in the bulk lipid region provides further evidence of the extent of membrane distortion caused by the protein (Figure S2).

Root mean square deviation (RMSD) and root mean square fluctuations (RMSF) revealed LpxR to be stable within the outer membrane, both in the *apo* form and when in complex with Re LPS. The RMSD of the barrel was 0.1-0.15 nm for both bound and *apo* forms of the protein, these values are similar to those reported from other simulation studies of outer membrane proteins^9^ (Figure S3). There was a marked decrease in RMSD of the alpha helical regions of the protein from 0. 3-0.35 nm to ~0.25 nm when comparing *apo* to bound forms of the protein.

The extracellular loops showed greater flexibility than the barrel in terms of RMSF, which agrees with other previously reported outer membrane proteins^10,11^; this is expected, as the extracellular residues are not afforded the same scaffold-like support as the barrel by bulk lipid tails. Details of consistency in tilt angle and system partial density between coarse-grain and united atom simulations, along with further information on RMSD and RMSF can be found in Figure S3.

### Conformational dynamics of the apo protein

Given the slow diffusion rate of LPS^12^, in order to study the conformational states of the protein, we decided to use a detergent micelle environment, which previously reported simulation studies been shown to allow for faster conformational dynamics ^13–15^. Over the course of three independent 500 ns simulations at 323 K (labelled ‘protein-in-micelle’ in Table 1), we observed substantial conformational rearrangements of the protein. While the β-barrel retained its original conformation as expected, the extracellular loops rearranged such that access to the LPS-binding pocket (Figure 2a and 2b) was occluded. Furthermore, residues previously identified as key for the catalytic process; N9, T34 and H122 were observe to form intermolecular hydrogen bonds. Thus a putative closed conformation of the protein has been identified from our simulations.

**Figure 2.**
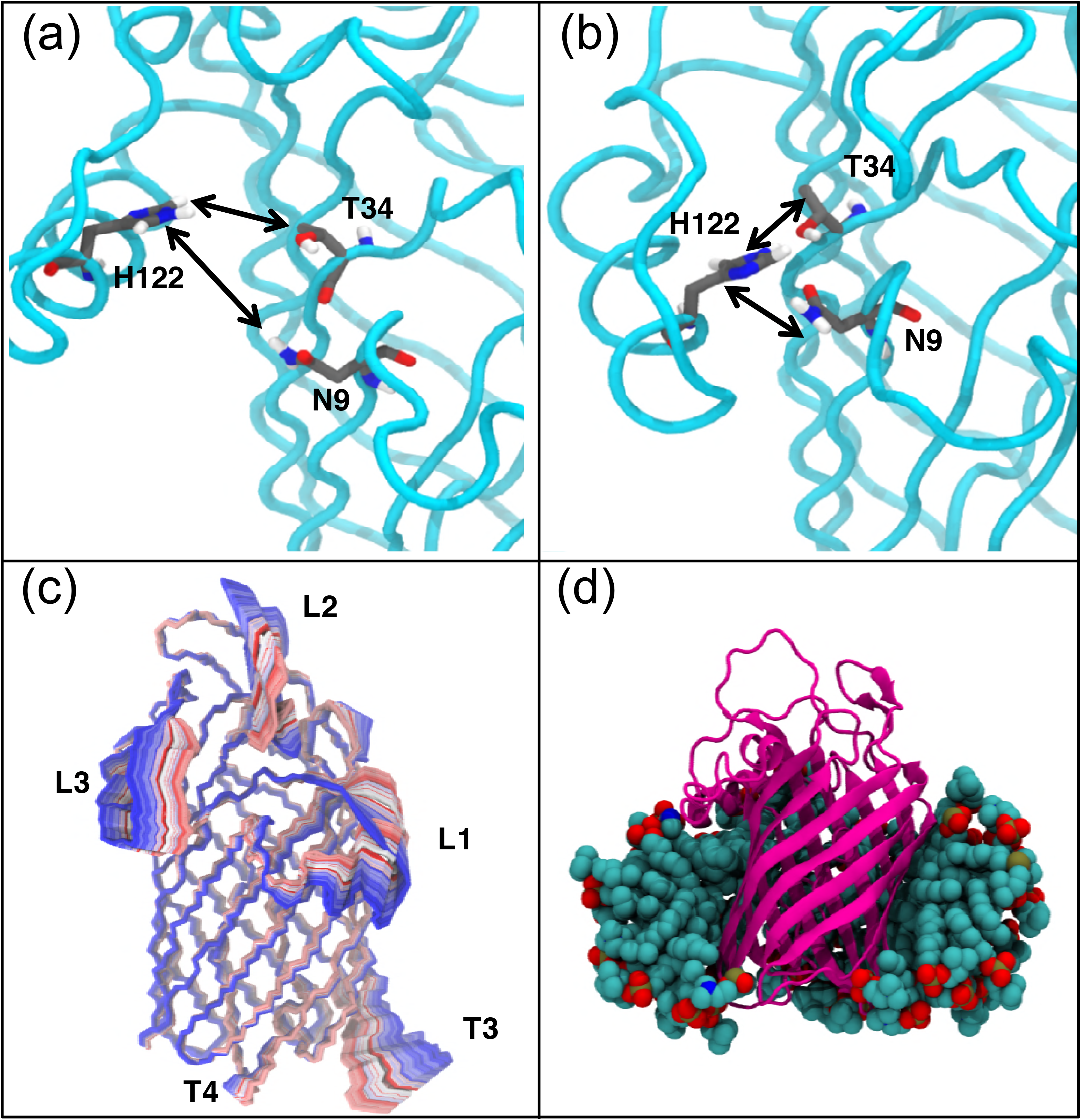
Conformational dynamics of LpxR within a detergent micelle at 323 K. (a) Position of catalytic residues in binding pocket in the energy minimised LpxR structure (PDB code 3FID). Protein backbone in cyan, with carbon, oxygen, nitrogen and hydrogen coloured grey, red, blue and white respectively. (b) Closed conformation of the protein revealed by simulations in DPC micelles, with catalytic residues within hydrogen bonding distance of each other. (c) The motion of the protein is depicted by extrapolating between the two extreme projections described by eigenvector 1 and then overlaying the conformation of the protein after every 20 ps. The image is coloured on a BWR scale (blue at the start of the simulation, through white, to red at the end of the simulation). (d) a representative snapshot of the protein in detergent micelle, some of the detergent molecules have been removed for clarity.

To further investigate the conformational space sampled by LpxR in a labile environment, we performed Principal Component Analysis (PCA) on the protein backbone by analysing a concatenated trajectory of all the micelle simulations. The first principal component accounted for 31.5% of the total variance in the backbone. The motion represented by this principal component was movement of extracellular loops L1, L2 and L3, which constitute the walls of the LPS-binding pocket to the ‘closed’ conformation of the protein (Figure 2c).

### Cation binding sites in the *apo* protein

Having established the structural stability of the protein, we then characterized the behaviour of the *apo* protein with respect to the putative Re LPS and cation-binding sites. We looked for evidence of a hydrogen bond between the carboxylate sidechain of E128 and Nε of H122, as is suggested in the proposed mechanism. However, across the 2.7 μs of *apo* protein in bilayer MD simulation, there was little evidence of a stable hydrogen bond here; due to steric hindrance produced by the backbone of extracellular loop L3 (residues 110-139, the α helix within this loop is defined by residues 110-126) it was easier for a hydrogen bond to form between the backbone carbonyl of E128 and Nε of H122. The distance between the centre of mass of H122 and E128 over 300 ns is shown in Figure S5a. We return to this later when discussing the mechanism of deacylation.

In each of the “model” simulations, the Ca^2+^ ion moved away from its initial binding site as soon as positional restraints were removed, indicating that either (i) a ligand is required to stabilize the ion within the binding site or (ii) this is not the cation binding site. When subject to positional restraints, the Ca^2+^ remained ~0.25 nm from T34; when restraints were removed, the ion moved away to ~1.2 nm from T34.

### Cation binding sites in the ligand-bound complex

The protein-ligand-membrane system was simulated in several states (Table 1); each one was initiated with the protein-lipid complex as predicted by docking calculations reported by Rutten et al^5^. In “model” simulations, the Ca^2+^ and water molecule proposed to be essential for the catalytic mechanism were retained, while in some simulations these were removed. We consider the former first. The Re LPS substrate remained non-covalently bound to the protein through electrostatic interactions between the Kdo sugar hydroxyl groups and the basic residues K67 and R68 for the duration of all of our simulations. Interestingly, Reynolds et al. noted that deep-rough LPS, which lacks Kdo sugars, is a poor substrate for LpxR(1). Our results suggest that this is likely due to the absence of the salt bridges between the Kdo sugars and the protein. Positioning of Re LPS with respect to the binding site on LpxR for “model” simulations may be seen in Figure 3a and 3c, and for “unbiased” simulations in Figure 3b and 3d; each image in Figure 3 was produced from a representative simulation at 323 K.

**Figure 3.**
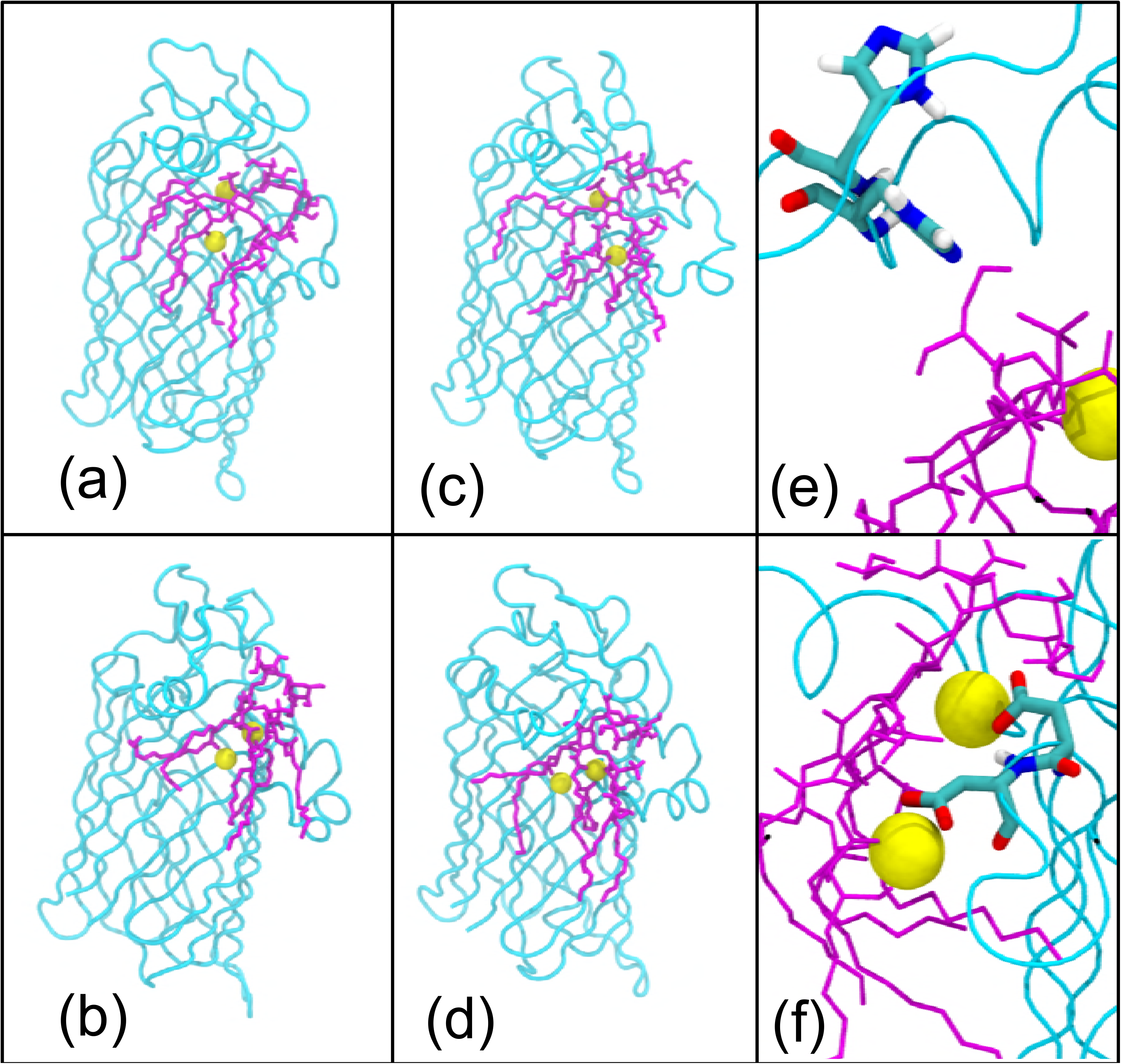
Snapshots of the protein-lipid-ion complex after 300 ns from two independent simulations of the ion biased (a) and (c) and ion unbiased (b) and (d) simulations. The two rotameric states of H122 are shown in panel (e) and panel (f) shows coordination of cations by residues D10 and D11. The protein is coloured cyan, Re LPS is magenta and ions are yellow. The membrane, water and other ions are omitted for clarity.

H122 sampled two major rotamers throughout the simulations (Figure 3e). The distance between the centre-of-mass of H122 and the scissile bond varied between 0.3 nm – 0.9 nm, although for most simulations the imidazole of H122 remained in a conformation angled toward the scissile ester bond. As expected, the H122A mutation did not affect protein interaction with Re LPS, nor did it affect the conformation of the L3 α helix (residues 110-126), as shown in Figure S5b.

Interestingly simulations of the wildtype protein predict a second cation-binding site in which an Mg^2+^ ion is coordinated by the glycerol oxygen adjacent to the scissile bond, the carboxylic acid moiety of D11 and water molecules found persistently in this region throughout the simulations. Coordination of cations by D10 and D11 is shown in Figure 3f. When the Ca^2+^ ion is removed prior to equilibration, the conformational behaviour of the putative active site is rather different. Although two Mg^2+^ ions are observed to enter the active site region during equilibration, and remain within this region throughout the simulations, neither one is located precisely in the same spot as the Ca^2+^ ion from the Rutten model(2). Furthermore, H122 is observed to flip out of the active site such that the side chain is pointing towards the extracellular loops. This movement of H122 is accompanied by snorkelling of K67 towards the phosphate group of the lipid A sugar, rather than interacting with the hydroxyl groups of the Kdo sugars as observed with the simulations of the Rutten model when Ca^2+^ is located within the proposed binding site. Residues T34 and D10 are not involved in interactions with the two cations in these simulations, instead one Mg^2+^ ion is observed to interact with D11 and two glycerol oxygens from adjacent acyl tails. The second cation-binding site identified in the wildtype simulations, is again observed here with, D11 involved in coordination along with a glycerol oxygen from one of the substrate lipid tails. The D11A mutation leads to the cation moving out of this binding site, indicating the importance of residue D11 for coordination of the second ion.

The scissile bond is oriented through interaction of the carbonyl oxygen with the Ca^2+^ ion, the latter is also coordinated by N9, D10, T34 and three water molecules throughout the simulations of the wildtype protein. In simulations of the protein with the single point mutation D10A, the Ca^2+^ moved out of the binding site such that after 300 ns it is no longer interacting with the protein, providing strong evidence that residue D10 plays a key role in coordination of the Ca^2+^ ion. In simulations of the T34A mutant however, the ion Ca^2+^ remains in the pocket, still coordinated by D10. Thus, our simulations suggest that for Ca^2+^ ion coordination, residue D10 is essential but T34 is not. Figure 4 shows the effect of specific residue mutation on the conformation of the protein-ion-lipid complex. Again, all images in Figure 4 were produced from simulations at 323 K.

**Figure 4.**
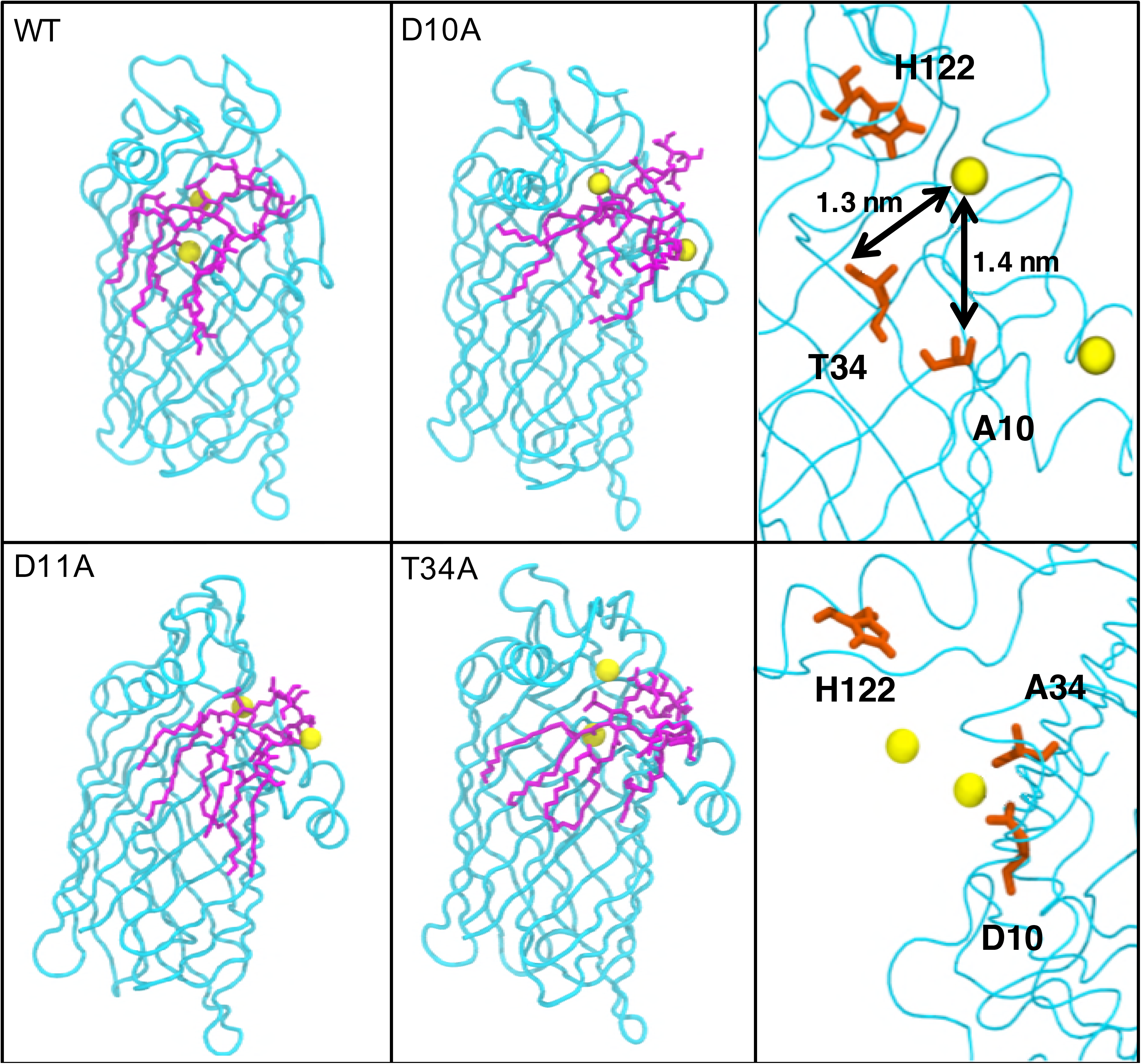
Snapshots of the protein-lipid-ion complex with specific residue mutations after 300 ns, compared to the wild-type. The membrane and solvent have been omitted for clarity. The top right panel shows the cation moving away from residue 10 when it is mutated from D to A. The bottom right panel shows the cation is still near its original location when residue 34 is mutated from T to A. The colour scheme is the same as Figure 3.

Our simulations therefore predict that both the aspartate residues within the putative binding site; D10 and D11 play a role in binding cations. Interestingly when Ca^2+^ is not coordinated by D10, the protein remains stable and the protein-substrate complex does not dissociate over the timeframe of our simulations, however the local conformational rearrangements of the nearby residues are such that it is difficult to envisage a deacylation process occurring. As such, our simulations support the hypothesis of Rutten et al. in which D10 must coordinate a cation. However, this does leave open the question of why does D11 also bind a cation? Cation binding in this region is observed in all our simulations, other than those of the D11A mutant, and therefore cation binding by D11 is likely to be important for ion recruitment to the active site.

### Catalytic mechanism

Having identified the conditions under which cations bind to the enzyme, we next sought to gain some insights into the catalytic mechanism of deacylation by LpxR by combining the results of MD and MC simulations, with clues from what is known about the mechanisms of phospholipase A2 enzymes^7^ and a previously hypothesised mechanism for deacylation from the static X-ray structure, docking studies and mutagenesis studies^5^. It is important to note here that initial predictions do not specify whether the mechanism requires one mechanistic water or a cascade of two water molecules, the latter is known to be the case for the mechanism of a secreted pancreatic phospholipase A2 enzyme^6,7^. Attention was paid to the number of water molecules found between H122 and the scissile ester during GCMC calculations. The putative catalytic protein residues, Ca^2+^ ion known to be essential for activity, and the ester moiety of the lipid substrate are shown in Figure 5.

**Figure 5.**
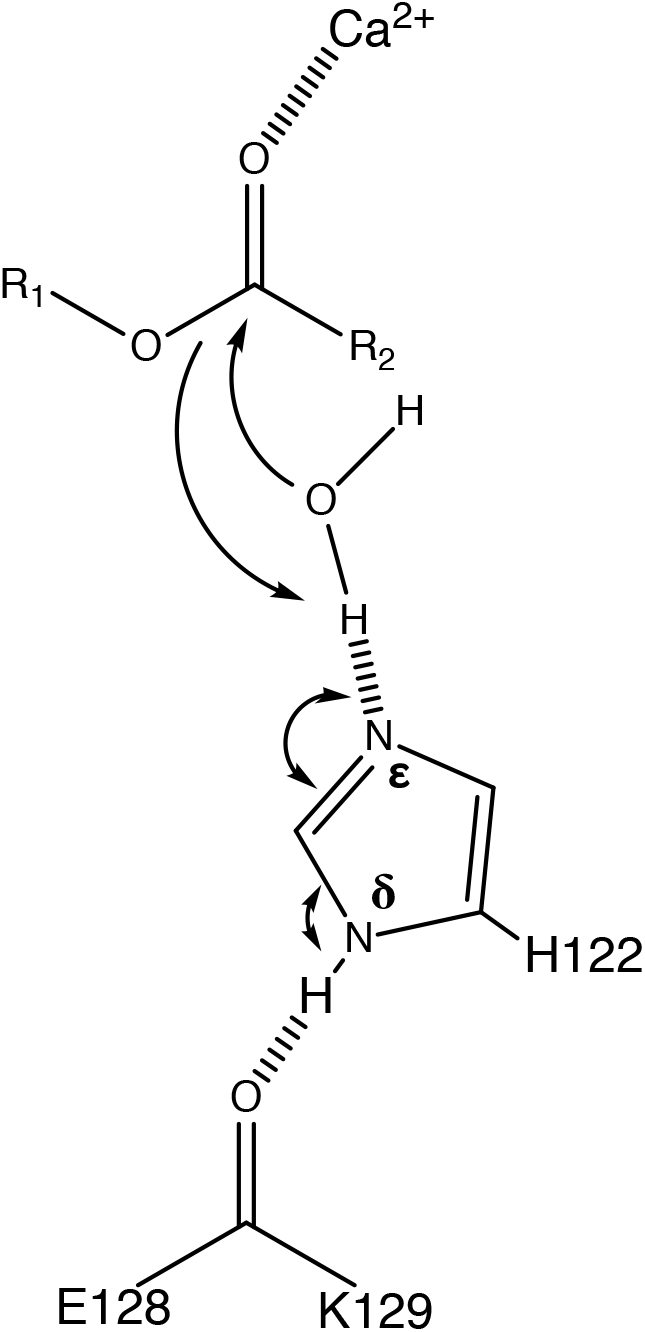
The residues E128, K129, H122, the portion of Re LPS to be cleaved and one Ca^2+^ ion.

To study aspects of the mechanism of deacylation and in particular to predict the number of water molecules likely to be involved in the mechanism, we performed GCMC simulations to determine the positions of water molecules near the protein-ligand complex. Simulations were performed at B_eq_, which corresponds to equilibrium with bulk water. The two protonation states of H122 were once again studied. H122 was more mobile in simulations in which it was Nδ-protonated. In two of the three independent simulations, the H122 shifted its position by rotating away from the Re LPS ester group, leaving a predominantly dry vacancy between H122 and the ligand. In the one independent repeat simulation in which the directional interaction between the H122 and the ester group was maintained, two water molecules were located by GCMC in-between the two moieties as part of a chain of water molecules. While the molecules were suitably positioned, the orientation of the water was such that formation of a hydrogen bond to the ester group of the substrate was not possible. However, the water was able to form a compensating hydrogen bond with another nearby water molecule. A snapshot of this water network and hydrogen bonding interactions is shown in Figure 6.

**Figure 6.**
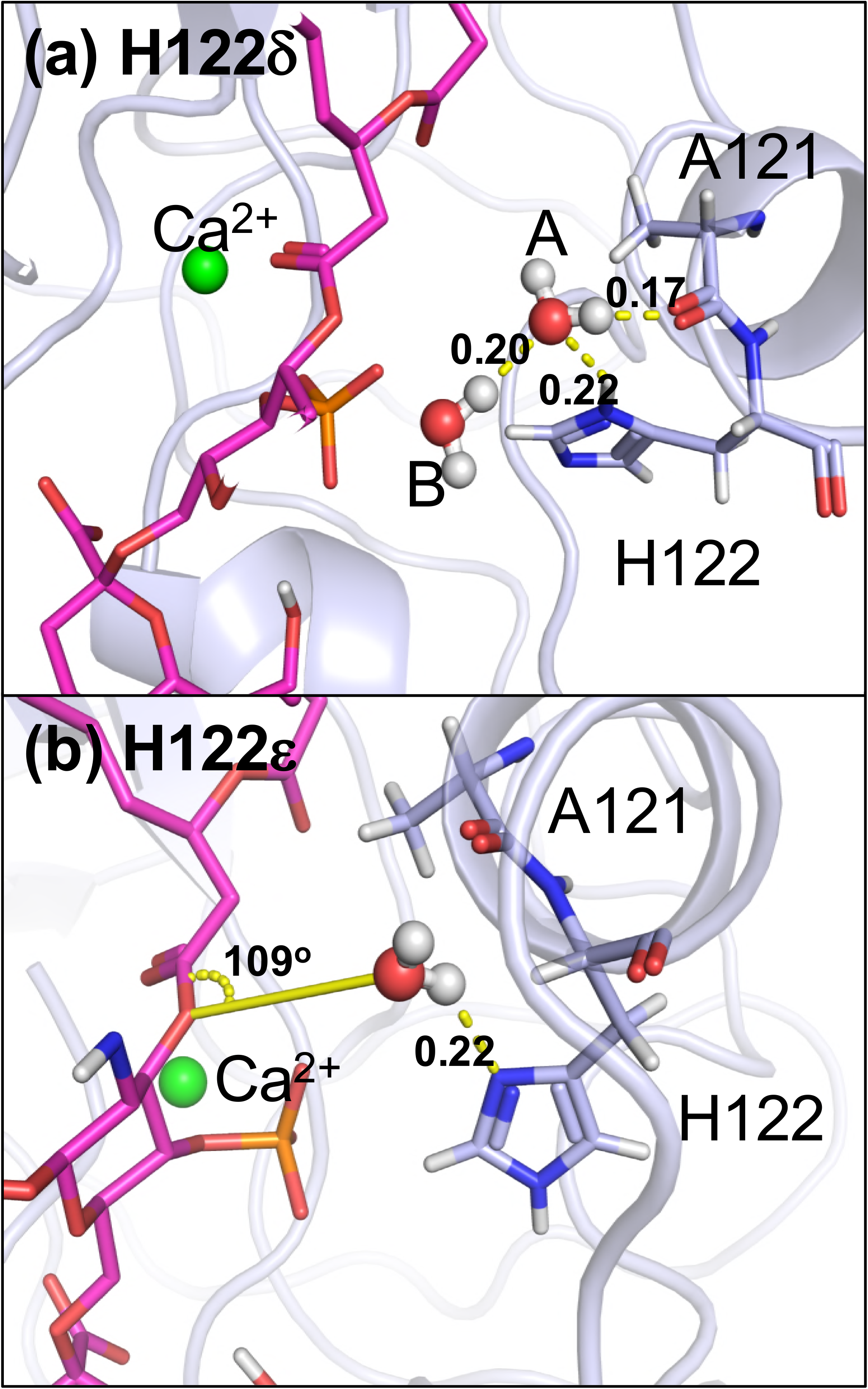
(a) Water locations from the GCMC simulation where H122 is Nδ-protonated. Two water molecules found between the H122 and the ester. Hydrogen bonding interactions are illustrated with a dashed yellow line, with length shown in nm. Water molecule B is forming hydrogen bonds with the H122 Hδ, water A and A121. The water A molecule is ~0.3 nm from the ester group throughout the simulation and is orientated with the oxygen atom pointed towards the group, which means that it is unable to hydrogen bond to the ester group and would therefore not result in hydrolysis. The water is stabilized in this orientation by hydrogen-bonding to another water close by. (b) Snapshot of the Nε-protonated H122. A121, H122, and ligand shown in stick representation, protein backbone shown in light blue. Calcium ion is shown in green. The relevant water(s) in each case are shown in red and white. Part of the ligand, and other GCMC water molecules have been removed from the image for clarity.

When the H122 was Nε-protonated, the hydrogen bonding interaction was flipped, with a hydrogen bond accepting Nδ oriented towards the ester bond. In all three repeats, the GCMC method located a single water molecule interacting with both the H122 and the ester bond. The orientation of this water molecule, shown (Figure 6) was such that it would enable it to act as a nucleophile in the catalysis. This water molecule was ~0.2 nm away from His122 (NδØto water 0) and ~0.3 nm away from the ester (water O to carbonyl C) throughout the simulation. The oxygen atom of the water molecule was at an angle of 109° to the plane of the ester group, close to the Bürgi-Dunitz angle of 107°^16^. Based on the clustering of the GCMC inserted water molecules, the water was present in this position 82.9% of the time when H122 was Nε-protonated. The binding free energy of the catalytic water determined using GCMC titration simulations for all combinations of rotameric and tautomeric states of H122, conformations of which are shown in Figure S6. The water molecule in the catalytic position was found to have binding free energies of; Nε -8.5 (0.2), Nδ -5.9 (0.3), Nε flipped -6.0 (0.2) and Nδ flipped -7.0 (0.2) kcal mol^-1^ respectively, with standard deviations shown. The corresponding GCMC titration curves are provided in Figure S7. In all cases the water molecule is tightly bound, but the binding free energy is most favourable for the Nε conformation – the proposed catalytic conformation. The orientation of this water molecule relative to H122 and the ester bond, as well as its energetic stability, provide compelling support for it being the catalytic water molecule. We note here that Bahnson suggests that the catalytic mechanism of phospholipase A2 involves water activation by a histidine that is Nε-protonated^7^. Based on the GCMC results favouring the Nε-protonated of H122, and the results from our molecular dynamics simulations, we can predict a mechanism for deacylation. We hypothesize here that the role of Ca^2+^ in the deacylation is to further polarise the sn2 carbonyl oxygen of the ester bond. The mechanism shown in Figure 7 has H122 directly increasing the nucleophilicity of the catalytic water via hydrogen bonding from Nδ. As previously mentioned, while E128 has been hypothesised to stabilize the orientation of H122 through hydrogen bonding, we do not observe a persistent E128-H122 hydrogen bond in any of our simulations, regardless of the protonation state of H122. The lack of hydrogen bonding between these two residues could in fact account for the relatively low activity of LpxR in *S. typhimurium*^5,17^; it remains to be seen whether a modified membrane composition could alter LpxR orientation and conformation, thereby stabilising the proposed E128-H122 hydrogen bond to increase the basicity of H122 and therefore activating the catalytic residue.

**Figure 7.**
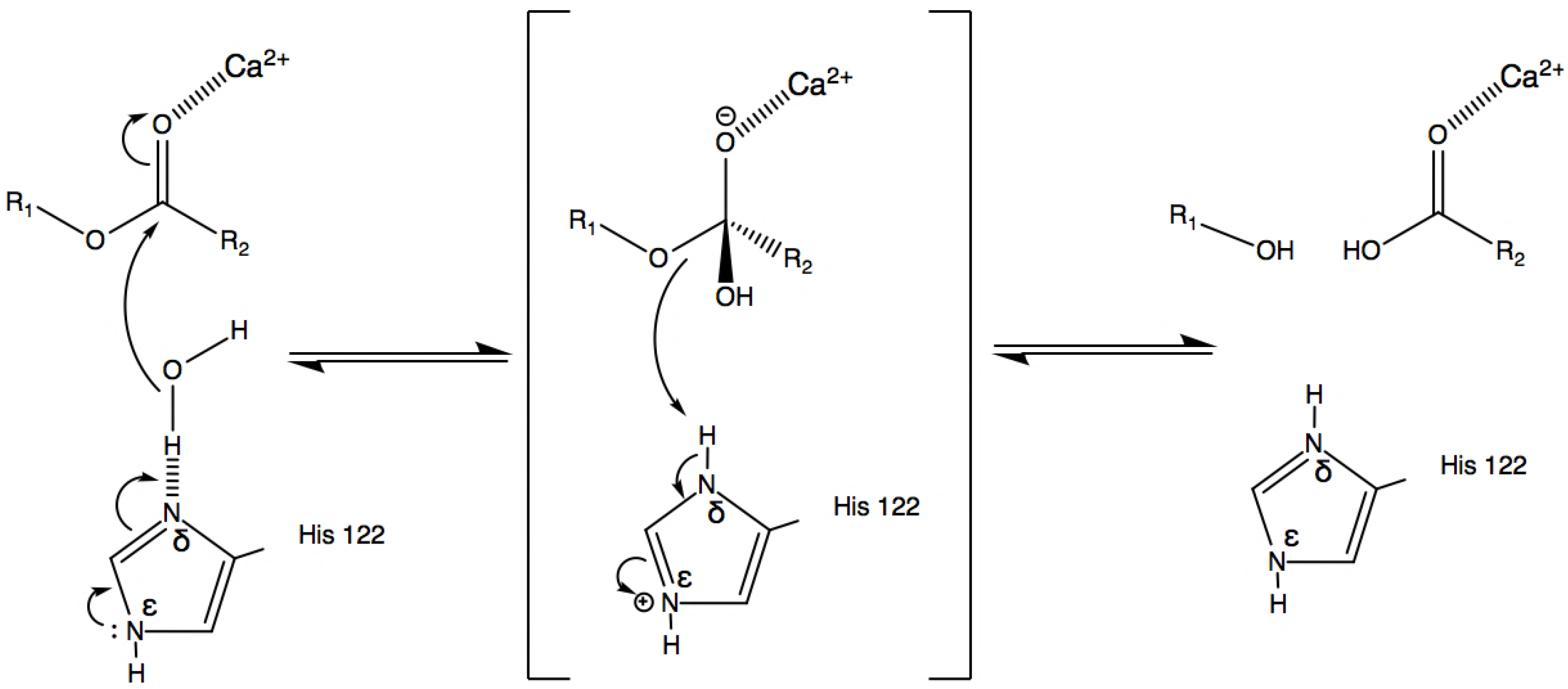
The predicted mechanism of deacylation based on our MD and GCMC simulations. The catalytic histidine, H122, is Nε-protonated.

### Protein and substrate after deacylation

To predict the diffusion of the products that occurs post-catalysis, the Re LPS ligand was manually deacylated by removing the covalent bond that would be enzymatically cleaved by LpxR. Simulations initiated from with the two post catalysis substrates within the hypothesised LPS binding region reveal that the cleaved acyl tails rapidly move away from the protein, towards the lower leaflet of the outer membrane. Meanwhile the modified LPS molecule remains within the putative binding site for the duration of our simulations. Each one of our six post-catalysis systems show the acyl tails moving towards the inner leaflet within 500 ns, as shown in Figure 8. Images in Figure 8 are produced from the bond break unbiased simulation at 323 K.

**Figure 8.**
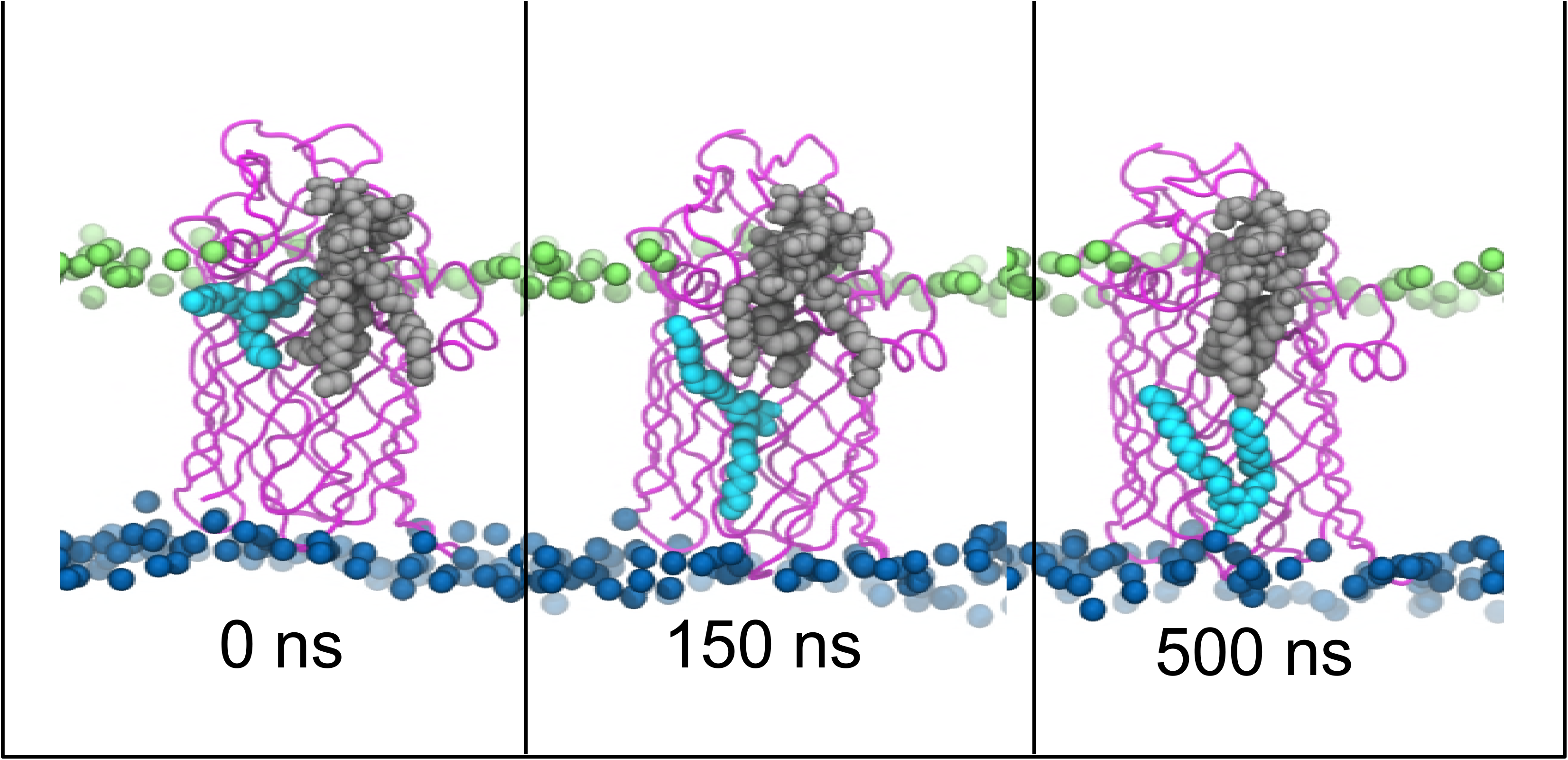
Movement of catalytically cleaved acyl tails (cyan) away from the protein (magenta) and the truncated Re LPS substrate (grey).

In the case of OmpLA, movement of lipid tails towards the inner leaflet leads to increased permeability and fluidity of the inner leaflet. This plays a role in bacteriocin release; it remains unclear as to whether LpxR could be implicated in the same way, or whether the acyl chains are promptly repurposed in lipid synthesis.

It is useful to consider the main limitation of the current study. This arises from the initial enzyme-substrate originating from a docked model. While the simulations show that lipid A is stable in the docked conformation within the putative binding site, structural data would confirm the structure of the enzyme-substrate complex. It is also worth noting that our study omits polarisation and other quantum effects, inclusion of which, which would allow for a more detailed study of the chemical reaction mechanism. Having said that, quantum mechanics calculations of membrane embedded proteins are challenging, with very little available in the literature in terms of best practice. The approach we have used here of studying multiple protonation states of the catalytically active residue, and using GCMC to identify water binding sites provides a sound classical alternative to using quantum methods for such a study.

The mechanism of deacylation by LpxR is of immense interest from an immunological and microbiological viewpoint, given that modification of LPS can lead to antimicrobial resistance. Here we have employed Molecular Dynamics and Monte Carlo simulations to study the conformational behaviour of the protein, which has enabled us to predict the structure of the putative ‘closed’ state of the protein in the absence of a substrate. The closed state of the protein exhibits hydrogen bonds between residues known to be key for the catalytic process. Thus we hypothesise that in the absence of substrate, these residues play a role in stabilising the protein in the closed conformation. For the substrate-bound state of the protein, we identify key protein-substrate interactions that hold the substrate within the active site. For the catalysis itself, we show that residue D10 plays a key role in coordinating a divalent cation. Our simulations predict the catalytic histidine, to be Nε-protonated, which differs from the previously proposed mechanism from structural data and docking studies in which the histidine would have to be Nδ-protonated. Furthermore, we provide quantitative evidence that one water molecule occupies the space between the catalytic histidine and the scissile bond in a tightly bound conformation, and thus seems suitably placed to participate in the hydrolysis reaction. We show that the tails that are removed from the LPS molecule are able to rapidly diffuse towards the inner leaflet and presumably insert into this leaflet, whereas movement of the remainder of the LPS molecule out of the active site, is slower. The combined data from the different types of simulations enable us to hypothesise the full mechanism of catalysis and also the movement of the newly formed chemical species, post catalysis. Indeed we show the importance of considering the conformational dynamics of membrane enzymes and ligands in a suitable local environment when attempting to decipher their mechanism of action.

## METHODOLOGY

### Simulation systems

The coordinates of the LpxR X-ray structure (pdb code 3FID), with Re-LPS, Ca^2+^ ion and a single water molecule modelled as described by Rutten at al., were obtained from Piet Gros^5^. The Ca^2+^ ion is positioned such that it replaces the zinc ion that was resolved in the X-ray structure, given that the former is known to be essential for catalytic activity. These initial coordinates were manipulated to set up a series of simulations that are summarised in Table 1

The protein, Ca^2+^ ion and single water molecule modelled by Rutten et al.^5^, were retained in all “model” simulations. The Re LPS molecule was removed from the putative binding site in “apo” simulations, and the Ca^2+^ ion and single water molecule were removed in “unbiased” simulations. In “Bond Break” systems, the scissile covalent bond of Re LPS was removed and the termini were protonated, thereby removing two acyl chains from the molecule. The two resultant molecules were parameterized in a manner consistent with the gromos54a7 force field^18^. Both molecules were retained in the simulations. Once the LPS molecule had been converted into two smaller molecules, these systems were then subjected to an additional 500 ns of production simulation. Table 1 summarises the four mutant proteins also studied, they were constructed by mutating the residues of interest using the PyMOL code^19^. In “H122_p” simulations, H122 was Nε-protonated, rather than Nδ-protonated as in all other simulations.

The model membrane used in each simulation was a mix of 90% phosphatidylethanolamine (PE), 5% phosphatidylglycerol (PG) and 5% cardiolipin in the inner leaflet, and Ra LPS in the outer leaflet. This is the same membrane composition previously used in the study of OmpA, a comparable outer membrane, β-barrel protein to LpxR^20^. As in the previous study, Mg^2+^ ions were used to neutralise the system.

### Coarse-grain Simulation Protocols

Coarse-grain simulations of the protein in an Ra LPS-phospholipid bilayer were performed to determine the preferred orientation of the protein within the membrane. Coarse-grain simulations used the MARTINI force field, with the LPS parameters of Hsu et al^21^. The Parrinello-Rahman barostat was used for semiisotropic pressure coupling with a time constant of 1 ps. The velocity rescale thermostat was used, with a time constant of 1 ps. The time step for integrations was 10 fs. Coulombic interactions were cut off at 1.4 nm and van der Waals reduced to zero between 0.9 – 1.4 nm. GROMACS molecular dynamics software package (version 5.1.4)^22,23^.

### Atomistic Simulation Protocols

Simulations were set up and performed using the GROMACS molecular dynamics software package (version 5.1.4) with the gromos54a7 force field^18,22,23^. The parameters for Re LPS and Ra LPS are identical to those described by Piggot et al. and Samsudin et al., respectively^12,20^. The SPC water model was used throughout the simulations^24^. Systems were maintained at temperatures of either the biological 310 K, or slightly higher at 323 K to improve sampling, using the Nosé-Hoover thermostat with a time constant of 0.5 ps^25,26^. The pressure of the system was maintained at 1 atm, with a time constant of 5 ps, using semi-isotropic pressure coupling with the Parrinello-Rahman barostat^27,28^. All van der Waals interactions were cut off at 1.4 nm and a smooth mesh Ewald (PME) algorithm was used to treat electrostatic interactions with a short-range cut off of 1.4 nm. Simulation parameters were chosen based on similar published studies of OmpA^20^. Each system was subjected to 500 ps of NVT simulation, followed by 20 ns of NPT for equilibration purposes. Positional restraints (1000 kJ mol^-1^ nm^2^) were placed on the Cα atoms of the protein and modelled Ca^2+^ ion and water molecule during NVT and NPT equilibration. For some of the simulations, these are highlighted in Table 1, the restraints were kept in place for the first 100 ns of the production runs. Production runs of 300 ns or 500 ns were then performed. The results were analysed using GROMACS^22,23^ tools and in-house scripts. Visualisation was performed using the VMD software package^29^.

### Micelle Simulations

To study protein structure in a more labile environment, LpxR was placed in a box with 100 dodecylphosphocholine (DPC) molecules, Na+ ions to neutralise charge and SPC water. Using a detergent micelle to study protein structure is a well-established method both *in silico* and *in vivo*^13,30,31^. Positional restraints (1000 kJ mol^-1^ nm^2^) were placed on the Cα atoms of the protein during NPT equilibration, during which a DPC detergent micelle formed around the protein. DPC parameters were downloaded from http://wcm.ucalgary.ca/tieleman/downloads. Equilibration lasted for 50 ns at 350 K to ensure the micelle remained localised around the protein. Production runs of 500 ns were then implemented at 323 K. Principal component and cluster analyses were performed on the resultant trajectories.

### Grand Canonical Monte Carlo Simulation Protocols

Grand canonical Monte Carlo (GCMC) simulations allow for the determination of both the location of water molecules within a defined region and their binding free energies. GCMC involves simulating in the grand canonical ensemble; that is the μVT ensemble, where μ is the chemical potential of the system. The chemical potential of the simulation controls the water occupancy of a GCMC region. A B value, proportional to the chemical potential will be used from this point forward, as it encapsulates both the chemical potential and additional constant parameters. A detailed explanation of these simulations, including a relation of B value and chemical potential have been published by Ross et al^32,33^. Two types of GCMC simulations were performed; one to calculate the locations of water molecules around the site of esterification using a larger GCMC box, and the other simulations to calculate the binding free energy of the water molecule identified as being likely to be catalytic using a smaller GCMC box.

Within one of the Ligand-Bound MD simulations at 100 ns, H122 was seen to move closer to the scissile bond, from ~0.9 nm to 0.6 nm. A snapshot of the system was taken from this and used as the starting point for Grand Canonical Monte Carlo (GCMC) simulations using ProtoMS^34^. The membrane was discarded and the protein-ligand complex solvated in a 4.5 nm sphere of TIP4P water^35^. The membrane was discarded to simplify calculations – as the GCMC box was to be placed over an area accessible by bulk solvent, it was unnecessary to include additional lipids. The protein and ligand were modelled using the Amber 14SB and gaff16 force-fields respectively^36–38^. A Ca^2+^ ion (parameters taken from^39^) was included in the simulation, where the location of the ion was taken from our atomistic trajectories. Given the histidine is of catalytic importance, to identify the correct protonation state and provide clues as to its mechanistic role, two sets of simulations were performed; one in which the Nδ was protonated and the other in which Nε was protonated. Due to the similar electron density of carbon and nitrogen, they are difficult to differentiate between in crystallographic electron density. This means that it can be difficult to resolve the orientation of histidine residues experimentally. For this reason, the alternative rotameric states (referred to as the *flipped* forms) of both the Nδ and Nε will be considered in the binding free energy simulations. For all simulations, 1 million (M) GCMC only equilibration steps, 1 M GCMC with protein ligand sampling equilibration steps were performed. The protein was sampled as fully flexible with both backbone and side chain moves. For the larger GCMC box, 40 M production steps were performed, while 20 M production steps were performed for the binding free energy calculations. For the binding free energy simulations, where multiple B values are simulated, replica exchange in B was attempted every 100,000 steps^33^. A list of all GCMC simulations can be found in Table 2. Location simulations: A large GCMC box of dimensions 0.73 x 0.49×1.16 nm^3^ was placed over the region of mechanistic interest – H122 and the Re LPS ester group. This simulation was performed to determine the hydration sites over the region. GCMC was performed at the equilibrium *B* value, -7.94, following the ProtoMS default methodology^34^. Three independent repeats were performed for both of the H122 tautomeric states.

Binding free energy simulations: In addition, GCMC calculations were performed to calculate the binding free energy of the specifically identified likely catalytic water molecule. The GCMC titration calculations were performed over a range of chemical potentials (16 equally spaced B values over an inclusive range of -40.79 to -10.79) for Nδ, Nε, Nδ flipped and Nε flipped H122 conformations. The water site of catalytic interest was defined as the water position found for the ε-protonated H122 of the larger GCMC box simulation. A small cubic GCMC box of length 0.2 nm was used, with the centre of the box was taken as the centroid of the water molecule. Three repeats were performed, and the binding free energy is calculated by performing 100 bootstrapping calculations of the data.

## Author contributions

G. M. S and H. B M performed the simulations. G. M. S, H. B M, J W. E and SK analysed the data. G. M. S, H. B M, and S. K. wrote the manuscript.

## References

1. Nikaido H. Molecular basis of bacterial outer membrane permeability revisited. Microbiol Mol Biol Rev. 2003;67(4):593–656.

2. Kim S, Patel DS, Park S, Slusky J, Klauda JB, Widmalm G, et al. Bilayer Properties of Lipid A from Various Gram-Negative Bacteria. Biophys J. 2016;111(8): 1750–60.

3. Hwang PM, Kay LE. Solution structure and dynamics of integral membrane proteins by NMR: a case study involving the enzyme PagP. Methods Enzym. 2005;394:335–50.

4. Rutten L, Geurtsen J, Lambert W, Smolenaers JJM, Bonvin AM, de Haan A, et al. Crystal structure and catalytic mechanism of the LPS 3-O-deacylase PagL from Pseudomonas aeruginosa. Proc Natl Acad Sci. 2006;103(18):7071–6.

5. Rutten L, Mannie J-PBA, Stead CM, Raetz CRH, Reynolds CM, Bonvin AMJJ, et al. Active-site architecture and catalytic mechanism of the lipid A deacylase LpxR of *Salmonella typhimurium*. Proc Natl Acad Sci U S A. 2009;106(6): 1960–4.

6. Berg OG, Gelb MH, Tsai MD, Jain MK. Interfacial enzymology: The secreted phospholipase A2-paradigm. Chem Rev. 2001;101(9):2613–53.

7. Bahnson BJ. Structure, function and interfacial allosterism in phospholipase A2: Insight from the anion-assisted dimer. Vol. 433, Archives of Biochemistry and Biophysics. 2005. p. 96–106.

8. Koebnik R, Locher KP, Van Gelder P. Structure and function of bacterial outer membrane proteins: barrels in a nutshell. Mol Microbiol. 2000;37(2):239–53.

9. Piggot TJ, Holdbrook DA, Khalid S. Conformational dynamics and membrane interactions of the E. coli outer membrane protein FecA: A molecular dynamics simulation study. Biochim Biophys Acta – Biomembr. 2013;1828(2):284–93.

10. Khalid S, Sansom MSP. Molecular dynamics simulations of a bacterial autotransporter: NalP from Neisseria meningitidis. Mol Membr Biol. 2006;23(6):499–508.

11. Ortiz-Suarez ML, Samsudin F, Piggot TJ, Bond PJ, Khalid S. Full-Length OmpA: Structure, Function, and Membrane Interactions Predicted by Molecular Dynamics Simulations. Biophys J. 2016;111(8):1692–702.

12. Piggot TJ, Holdbrook D a, Khalid S. Electroporation of the E. coli and S. Aureus membranes: molecular dynamics simulations of complex bacterial membranes. J Phys Chem B. 2011;115(45):13381–8.

13. Bond PJ, Sansom MSP. Membrane protein dynamics versus environment: Simulations of OmpA in a micelle and in a bilayer. J Mol Biol. 2003;329(5): 1035–53.

14. Bond PJ, Faraldo-Gomez JD, Deol SS, Sansom MSP. Membrane protein dynamics and detergent interactions within a crystal: A simulation study of OmpA. Proc Natl Acad Sci. 2006;103(25):9518–23.

15. Cox K, Bond PJ, Grottesi A, Baaden M, Sansom MSP. Outer membrane proteins: Comparing X-ray and NMR structures by MD simulations in lipid bilayers. Eur Biophys J. 2008;37(2):131–41.

16. B:urgi HB, Dunitz JD, Lehn JM, Wipff G. Stereochemistry of reaction paths at carbonyl centres. Tetrahedron. 1974;30(12):1563–72.

17. Reynolds CM, Ribeiro AA, McGrath SC, Cotter RJ, Raetz CRH, Trent MS. An outer membrane enzyme encoded by Salmonella typhimurium LpxR that removes the 3’O-acyloxyacyl moiety of lipid A. J Biol Chem. 2006;281(31):21974–87.

18. Huang W, Lin Z, Van Gunsteren WF. Validation of the GROMOS 54A7 force field with respect to β-peptide folding. J Chem Theory Comput. 2011;7(5):1237–43.

19. DeLano WL. The PyMOL Molecular Graphics System, Version 1.8. Schrödinger LLC. 2014;http://www.pymol.org.

20. Samsudin F, Ortiz-Suarez ML, Piggot TJ, Bond PJ, Khalid S. OmpA: A Flexible Clamp for Bacterial Cell Wall Attachment. Structure. 2016;24(12):2227–35.

21. Hsu P-C, Jefferies D, Khalid S. Molecular Dynamics Simulations Predict the Pathways via Which Pristine Fullerenes Penetrate Bacterial Membranes. J Phys Chem B. 2016;120(43):11170–9.

22. Van Der Spoel D, Lindahl E, Hess B, Groenhof G, Mark AE, Berendsen HJC. GROMACS: Fast, flexible, and free. Vol. 26, Journal of Computational Chemistry. 2005. p. 1701–18.

23. Martins-Costa MTC, Ruiz-López MF. Gromacs 5.1 manual. J Comput Chem. 2017;38(10):659–68.

24. Berendsen HJC, Postma JPM, Gunsteren WF Van, Hermans J. Interaction models for water in relation to protein hydration. Intermol Forces. 1981;331–42.

25. Nosé S. A molecular dynamics method for simulations in the canonical ensemble. Mol Phys. 1984;52(2):255–68.

26. Hoover WG. Canonical dynamics: Equilibrium phase-space distributions. Phys Rev A. 1985;31(3):1695–7.

27. Parrinello M, Rahman A. Strain fluctuations and elastic constants. J Chem Phys. 1982;76(5):2662–6.

28. Parrinello M, Rahman A. Crystal structure and pair potentials: A molecular-dynamics study. Phys Rev Lett. 1980;45(14):1196–9.

29. Humphrey W, Dalke A, Schulten K. VMD: Visual molecular dynamics. J Mol Graph. 1996;14(1):33–8.

30. Marrink SJ, Tieleman DP, Mark AE. Molecular Dynamics Simulation of the Kinetics of Spontaneous Micelle Formation. J Phys Chem B. 2000;104(51):12165–73.

31. Zhang L, DeHaven RN, Goodman M. NMR and modeling studies of a synthetic extracellular loop II of the κ opioid receptor in a DPC micelle. Biochemistry. 2002;41(1):61–8.

32. Ross GA, Bodnarchuk MS, Essex JW. Water Sites, Networks, and Free Energies with Grand Canonical Monte Carlo. J Am Chem Soc. 2015;137(47):14930–43.

33. Ross GA, Bruce Macdonald HE, Cave-Ayland C, Cabedo Martinez AI, Essex JW. Replica exchange and standard state binding free energies with grand canonical Monte Carlo. J Chem Theory Comput. 2017;acs.jctc.7b00738.

34. Bodnarchuk MS, Bradshaw R, Cave-Ayland C, Genheden S, Cabedo Martinez AI, Michel J, et al. ProtoMS [Internet]. Southampton: University of Southampton; 2017. Available from: http://www.essexgroup.soton.ac.uk/ProtoMS/

35. Neumann M. Dielectric relaxation in water. Computer simulations with the TIP4P potential. J Chem Phys. 1986;85(3):1567–80.

36. Case DA, Cheatham TE, Darden T, Gohlke H, Luo R, Merz KM, et al. The Amber biomolecular simulation programs. Vol. 26, Journal of Computational Chemistry. 2005. p. 1668–88.

37. Dickson CJ, Rosso L, Betz RM, Walker RC, Gould IR. GAFFlipid: a General Amber Force Field for the accurate molecular dynamics simulation of phospholipid. Soft Matter. 2012;8(37):9617.

38. Maier JA, Martinez C, Kasavajhala K, Wickstrom L, Hauser KE, Simmerling C. ff14SB: Improving the Accuracy of Protein Side Chain and Backbone Parameters from ff99SB. J Chem Theory Comput. 2015;11(8):3696–713.

39. Åqvist J. Ion-water interaction potentials derived from free energy perturbation simulations. J Phys Chem. 1990;94(21):8021–4.

